# Mechanism of CFTR correction by type I folding correctors

**DOI:** 10.1101/2021.06.18.449063

**Authors:** Karol Fiedorczuk, Jue Chen

## Abstract

Small molecule chaperones have been exploited as therapeutics for the hundreds of diseases caused by protein misfolding. The most successful examples are the CFTR correctors, which transformed cystic fibrosis therapy. These molecules revert folding defects of the ΔF508 mutant and are widely used to treat patients. However, their mechanism of action is unknown. Here we present cryo-electron microscopy structures of CFTR in complex with two FDA-approved correctors: lumacaftor and tezacaftor. Both drugs insert into a hydrophobic pocket in the first transmembrane domain (TMD1), linking together four helices that are thermodynamically unstable. Mutating residues at the binding site rendered ΔF508-CFTR insensitive to lumacaftor and tezacaftor, underscoring the functional significance of the structural discovery. These results support a mechanism in which the correctors stabilize TMD1 at an early stage of biogenesis, prevent its pre-mature degradation, and thereby allosterically rescue a large number of disease-causing mutations.

## INTRODUCTION

Cystic fibrosis is a genetic disease caused by mutations in the cystic fibrosis transmembrane regulator (CFTR), an anion channel that regulates salt, fluid, and pH balance in many organs (Cutting, 2015). CFTR belongs to the ATP-binding cassette (ABC) transporter family. It is a single polypeptide composed of two pseudo-symmetrical halves connected by a regulatory domain (RD). Each half of CFTR contains a transmembrane domain (TMD) that forms the ion conduction pathway and a cytoplasmic nucleotide-binding domain (NBD) that binds ATP. The RD, unique in CFTR, must be phosphorylated for the channel to open (Cheng et al., 1991). The phosphorylated CFTR channel is gated by ATP binding and hydrolysis (Csanády et al., 2019). The molecular structure of CFTR has been determined in two conformational states (Liu et al., 2017; Zhang et al., 2018). In the un-phosphorylated, ATP-free state, the RD lies in between the two NBDs and the pore is closed (Liu et al., 2017). The structure of the phosphorylated, ATP-bound state was obtained from the hydrolysis-deficient mutant E1371Q, which shows that phosphorylation releases the RD from its inhibitory position, permitting NBD dimerization and channel opening (Zhang et al., 2018).

More than 300 mutations lead to cystic fibrosis: some cause defects in channel function and others interfere with CFTR expression and folding (Welsh and Smith, 1993). The most common mutation is the deletion of phenylalanine at position 508 (ΔF508), which results in the retention of folding intermediates in the endoplasmic reticulum (ER) (Cheng et al., 1990). In recent years, CFTR modulators have been developed to revert the effects of the disease-causing mutations (Habib et al., 2019). Small molecules that enhance channel activity are called potentiators, and chaperones that increase the amount of folded CFTR are called correctors. Currently, one potentiator (ivacaftor) and three correctors (lumacaftor, tezacaftor, and elexacaftor) are in clinical use. Patients with folding mutations are treated with a combination therapy of potentiator and corrector.

Extensive research has been devoted to uncovering the mechanisms of CFTR modulators. Whereas the structural and functional basis of potentiator ivacaftor action has been described (Eckford et al., 2012; Jih and Hwang, 2013; Liu et al., 2019; Van Goor et al., 2009), the mechanism of correctors remains largely undefined. In one study it was suggested that lumacaftor (formally VX-809) acts through perturbation of membranes as it appeared to be homogeneously distributed throughout the lipid bilayer (Baroni et al., 2014). In multiple other studies, a direct action on the channel was proposed, but the location of the binding site remains in dispute (Eckford et al., 2014; Farinha and Canato, 2017; He et al., 2013; Hudson et al., 2017; Krainer et al., 2020; Krainer et al., 2018; Laselva et al., 2018; Loo et al., 2013; Loo and Clarke, 2017; Okiyoneda et al., 2013; Ren et al., 2013; Sinha et al., 2015). In this study, we determined cryo-EM structures of CFTR in complex with either lumacaftor or tezacaftor (formally VX-661). These structures identify the CFTR drug-binding site for these correctors and support a specific mechanism of action.

## RESULTS

### Lumacaftor and tezacaftor bind directly to CFTR

CFTR correctors have been categorized into different clusters based on their functional redundancy or additivity. Correctors in the same cluster do not exhibit additive effects and are proposed to share a similar mechanism (Veit et al., 2020; Veit et al., 2018). Correctors from different clusters act through different mechanisms and some can be combined to synergistically promote CFTR folding (Farinha et al., 2013; Pedemonte et al., 2005; Van Goor et al., 2011; Veit et al., 2020; Veit et al., 2018). Based on this categorization, lumacaftor (VX-809) and tezacaftor (VX-661) belong to the same cluster called type I correctors. They are structural analogues, both containing a 1,3-**B**enzodioxol-5-yl-**C**yclopropane **C**arboxamide (BCC) headgroup (Figure 1A). Other type I correctors such as C18 (Okiyoneda et al., 2013), ABBV/GLPG-2222 (Wang et al., 2018) (Figure 1A), and ARN23765 (Pedemonte et al., 2020) also share a similar chemical structure and likely a similar mechanism of action.

**Figure 1.**
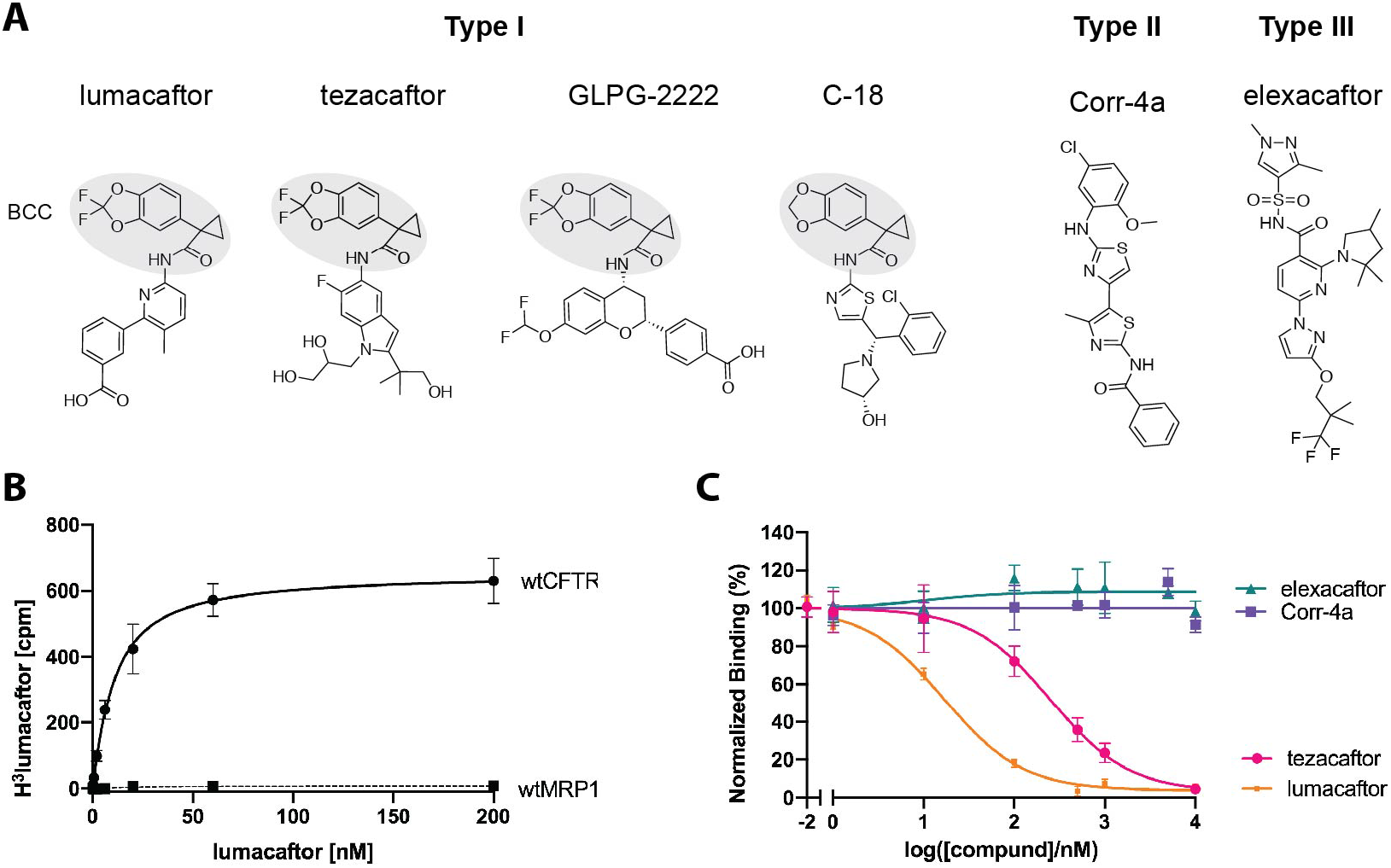
Lumacaftor and tezacaftor bind to CFTR competitively. **(A)** The chemical structures of representative type I, II and III correctors. The 1,3-**B**enzodioxol-5-yl-**C**yclopropane **C**arboxamide (BCC) headgroup is highlighted in grey. **(B)** Saturation binding and nonlinear regression analysis of [^3^H]lumacaftor binding to *wt*CFTR in the absence of phosphorylation and ATP (*K_d_* = 8.3 ± 2.2 nM). Also shown is a negative control using a related ABC transporter MRP1. **(C)** Competition binding assay. The binding of [H^3^]lumacaftor (10 nM) was plotted as a function of the competitor’s concentration. Data were fit to a single-site competitive binding model. The K_i_ values for lumacaftor and tezacaftor are 7.7 ± 2.0 nM and 0.12 ± 0.04 μM, respectively. No competition was observed for elexacaftor and Corr-4a. Each data point represents the mean and the standard error of the mean (SEM) of 3-9 of measurements.

Although lumacaftor was identified on the basis of its ability to increase cell surface expression of ΔF508-CFTR, its effect is not limited to this particular mutation. Lumacaftor promotes folding of many other mutations as well as the *wt*CFTR (He et al., 2013; Lukacs and Verkman, 2012; Moniz et al., 2013; Ren et al., 2013; Van Goor et al., 2011). A key question to address then is: Does lumacaftor act on misfolded CFTR to restore the tertiary structure or does it stabilize already-folded CFTR in its native conformation? The latter possibility is supported by two recent studies, which showed that lumacaftor, and its analog C18, bind and stabilize ΔF508-CFTR after its rescue to the cell surface (Eckford et al., 2014; Okiyoneda et al., 2013). C18 also binds directly to purified *wt*CFTR reconstituted in proteoliposomes (Eckford et al., 2014). To measure quantitatively the interactions between lumacaftor and purified *wt*CFTR, we established a scintillation proximity assay (SPA) showing that specific binding of [^3^H]lumacaftor increases as a function of its concentration (Figure 1B). The data fit well to a single-site binding model via nonlinear regression analysis, resulting in an equilibrium dissociation constant (K*_d_*) of 8.3 ± 2.2 nM. In comparison, the effective half concentration (EC_50_) of lumacaftor to rescue ΔF508-CFTR function is reported to be 81 ± 19 nM (Van Goor et al., 2011). The approximate 10-fold difference in the affinity/effective dose is likely explicable on the basis that the former measures the interaction of the drug with *wt*CFTR *in vitro* whereas the latter measures the cellular effects on the ΔF508 mutant.

To test if lumacaftor and tezacaftor share a common binding site, we performed a competition assay by measuring lumacaftor binding in the presence of increasing concentrations of tezacaftor (Figure 1C). Tezacaftor displaced [^3^H]lumacaftor in a manner quantitatively consistent with a 1:1 competitive mechanism with an inhibition constant (K*_i_*) equal to 115 ± 42 nM (Figure 1C), also comparable to the *in vivo* potency of tezacaftor (EC_50_ = 516 nM) (Van Goor et al., 2016; Van Goor et al., 2011). Using the same competition assay, the K*_i_* of unlabeled lumacaftor was determined to be 7.7 ± 2.0 nM (Figure 1C), consistent with the K_d_ value determined in the direct binding assay (Figure 1B). Two other structurally unrelated correctors, Corr-4a (Type II corrector) and elexacaftor (Type III corrector) (Figure 1A) did not displace [^3^H]lumacaftor (Figure 1C). The lack of competition by those two correctors is in agreement with folding studies showing that Corr-4a and elexacaftor function synergistically with lumacaftor (Pedemonte et al., 2005; Van Goor et al., 2011; Veit et al., 2020) and do not belong to the type I corrector cluster.

### Structural identification of the lumacaftor binding site

To identify the corrector-binding site, we determined the cryo-EM structures of the CFTR/lumacaftor complex in two conformational states (Figure 2A, S1, S2, S3, S4 and Table S1). In the absence of phosphorylation and ATP, *wt*CFTR exhibits an NBD-separated conformation as observed before (Liu et al., 2017; Zhang and Chen, 2016). The map, at an overall resolution of 3.9 Å, reveals a ligand density as strong as those of the protein mainchain atoms (Figure S7A). This density is absent from any of the CFTR maps obtained in the absence of a corrector (Figure S7B and S7D). The density has an elongated L shape consistent with the chemical structure of lumacaftor (Figure 1A). A higher resolution (2.7 Å) structure was determined from phosphorylated, ATP-bound CFTR(E1371Q), which exhibits the NBD-dimerized conformation as expected (Zhang et al., 2017, 2018). At the same ligand-binding location, we observe a similarly strong but better-defined density that fits lumacaftor unambiguously (Figure 2C-E and S7C). These results indicate that lumacaftor binds to CFTR in both conformational states.

**Figure 2.**
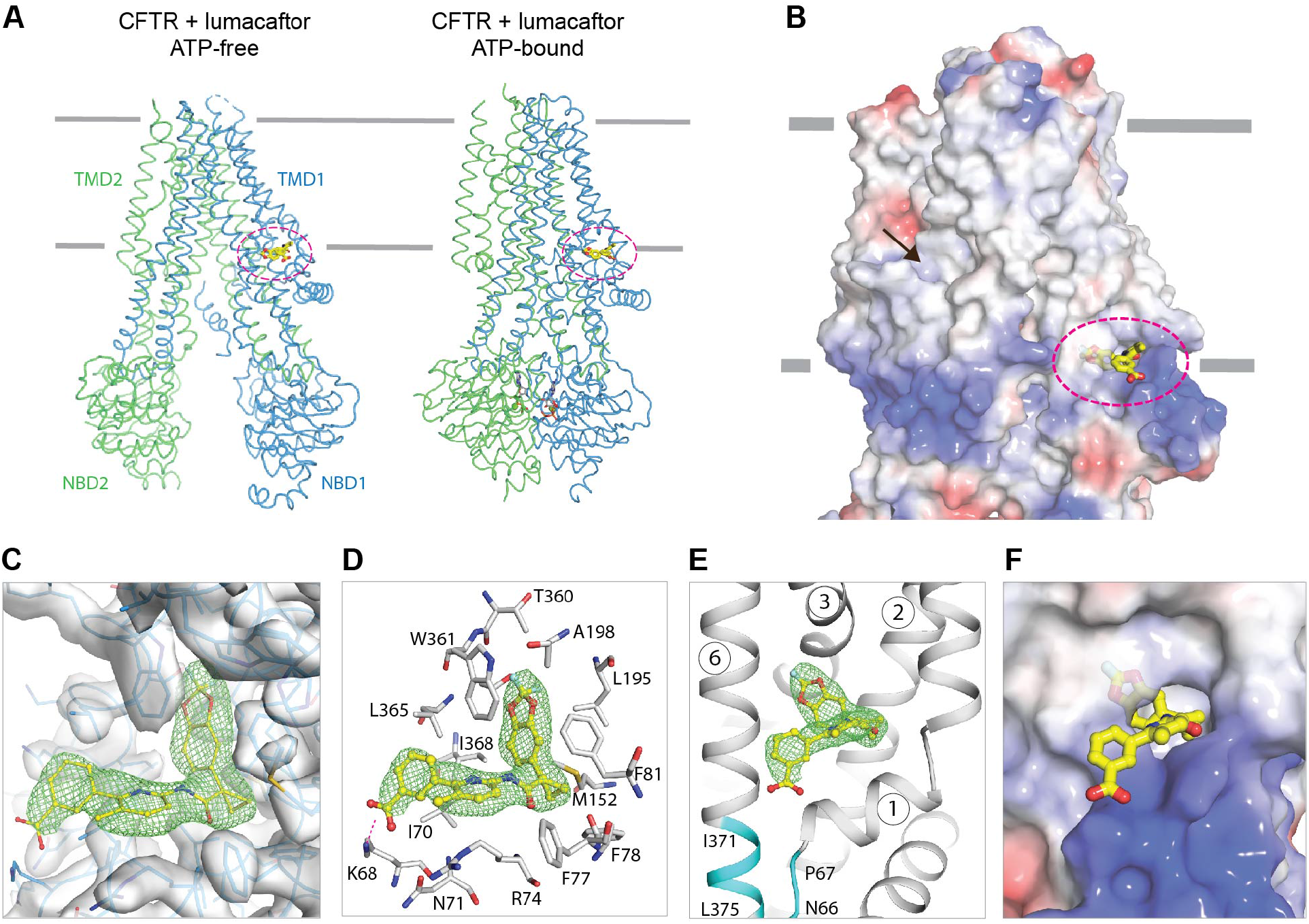
Lumacaftor binds to CFTR in both conformational states. **(A)** The overall structure of lumacaftor bound to the unphosphorylated, ATP-free *wt*CFTR (left) and phosphorylated, ATP-bound CFTR(E1371Q) (right). TMD1 and NBD1 are shown in blue, TMD2 and NBD2 in green. Lumacaftor is represented in yellow sticks and highlighted by the circle in magenta. The grey lines indicate the membrane boundaries. **(B)** Lumacaftor binds at the protein/membrane interface. The surface of the ATP-bound CFTR is shown by electrostatics and scaled from −10kT/e (red) to +10kT/e (blue). For reference, the location of the ivacaftor-binding site is indicated by an arrow. **(C)** Experimental density of the lumacaftor-binding site. Protein density is shown in grey and lumacaftor density in green. **(D)** Molecular recognition of lumacaftor. Residues within 4.5 Å of lumacaftor are shown as grey sticks. The salt bridge between K68 and lumacaftor is indicated by a magenta dashed line. **(E)** The lumacaftor-binding site is formed by TM 1, 2, 3, and 6. Cyan highlights the interactions between residues 371-375 and the N-terminal region of TM1. **(F)** Electrostatic surface representation of the same region as in Panel (E). See also Figures S1, S2, S3, S4, S7, S10, and Table S1.

The binding site is located in TMD1, at the level where the phospholipid head groups of the inner membrane meet the hydrophobic core (Figure 2A-B). Lumacaftor interacts with CFTR predominantly through van der Waals interactions, except for a salt bridge with K68 (Figure 2D and 3F). The BCC headgroup, a shared moiety among most type I correctors, inserts into a hydrophobic pocket formed by TM1, 2, 3, and 6 (Figure 2E-F). The shape of the BCC headgroup complements the narrow pocket in a classic “key in a lock” fashion (Berg et al., 2019). The polar half of lumacaftor extends outside the pocket, tethering the cytoplasmic ends of TM1 and TM6 together by interacting with residues 70-74 on TM1 and L365 and I368 on TM6 (Figure 2D-E).

**Figure 3.**
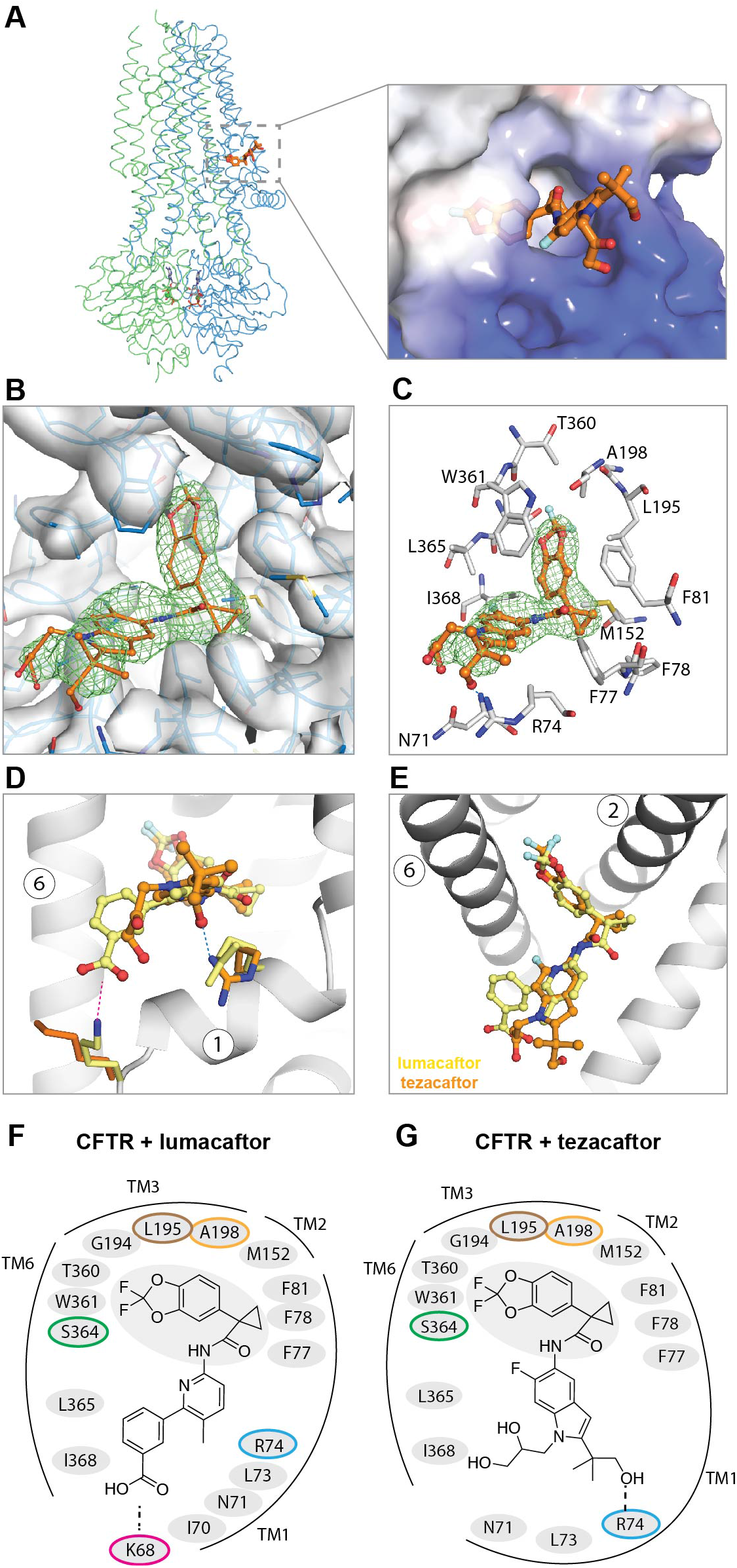
Tezacaftor binds CFTR at the same site as lumacaftor. **(A)** The overall structure of the CFTR/tezacaftor complex, with a zoom-in view of the binding site. Tezacaftor is represented in orange sticks and the protein surface is colored by electrostatics, scaled from −10kT/e (red) to +10kT/e (blue). **(B)** Experimental density of the tezacaftor-binding site. Protein density is represented in grey and tezacaftor density in green. **(C)** Molecular interaction at the binding site. Restudies within 4.5 Å distance from tezacaftor are shown as grey sticks. The H-bond between R74 and tezacaftor is indicated as a blue dashed line. **(D) (E)** Two views to compare the structures of lumacaftor (yellow) and tezacaftor (orange). The side chains of K68 and R74 are shown to highlight their different roles in drug binding. **(F) (G)** Schematic drawing of the CFTR-corrector interactions. All the restudies within 4.5 Å distance of the corrector are depicted. Residues mutated in the maturation and binding assays are indicated with colored circles. See also Figures S5, S6, S10, and Table S1.

In agreement with the structure, previous *in vivo* studies showed that the minimal domain sensitive to lumacaftor contains the N-terminal 375 residues (Ren et al., 2013). Removing residues 371-375 rendered lumacaftor ineffectual (Ren et al., 2013). Although residues 371-375 do not contact lumacaftor directly, they interact with N66 and P67, positioning TM1 to coordinate lumacaftor (Figure 2E, cyan ribbon). These observations underscore the structural role of residues 371-375 in constructing the binding site; they also support the previous conclusion that lumacaftor binds to TMD1 in its folded state (Eckford et al., 2014; Okiyoneda et al., 2013).

The location and the chemical nature of lumacaftor-binding are very different from those of the potentiator ivacaftor (Liu et al., 2019). Lumacaftor inserts into a deep pocket in TMD1 and its affinity is mediated through a high degree of shape complementarity, which maximizes van der Waal’s interactions. In contrast, ivacaftor binds to a shallow cleft on TMD2 at the center of the membrane (Figure 2B, arrow and S7D). Mutagenesis studies have shown that hydrogen bonds, rather than van der Waal’s interactions, play a predominant role in ivacaftor recognition (Liu et al., 2019).

### Tezacaftor binds to the same site as lumacaftor

The competition assay data suggested that lumacaftor and tezacaftor share an overlapping binding site on CFTR. To interrogate this conclusion, we determined the structure of tezacaftor-bound CFTR in the NBD-dimerized conformation to 3.8 Å resolution (Figure 3A-C, S3, S6 and Table S1). Indeed, a ligand density is observed at the same location in TMD1 (Figure 3B-E). Inside the hydrophobic pocket, the density also has an elongated shape that fits the BCC headgroup. The density outside the pocket is different than in the lumacaftor-bound structure in a manner consistent with structural differences between the two correctors (Figure 1A and 3B-C).

The BCC headgroup of tezacaftor interacts with the same set of pocket-lining residues as lumacaftor (Figure 3C and 3G). The polar region of tezacaftor, also exposed at the protein/lipid interface, interacts with CFTR in a slightly different manner. Instead of forming a salt bridge with K68, tezacaftor forms an H-bond with R74 (Figure 3C-G). In addition, tezacaftor interacts with fewer residues in TM1 (Figure 3G), which may explain the relatively lower affinity (Figure 1C) and lower potency of tezacaftor (Van Goor et al., 2016; Van Goor et al., 2011). Comparison of correctors-bound and -free structures reveals little difference, except for the dispositions of the K68 and R74 side chains. Upon drug binding, the terminal nitrogen on K68 moves about 4 Å to interact with lumacaftor and the side chain of R74 moves 3 Å to interact with either drug (Figure S8).

### Binding-site mutations reduce the efficacy of lumacaftor and tezacaftor in rescuing ΔF508-CFTR

Because the structures of drug complexes are of folded CFTR, one might ask whether the structurally identified binding site is the same site of action during CFTR biogenesis. To address this question, we introduced binding site mutations to the ΔF508-CFTR background and analyzed the ability of correctors to rescue these mutants (Figure 4A). Based on the structure, we reason that substituting small pocket-lining residues with larger ones would likely produce steric occlusion of both lumacaftor and tezacaftor. The effects of correctors were quantified using an established gel-shift assay, which measures the abundance of the fully glycosylated CFTR relative to the core-glycosylated form (Figures 4A and S9). As CFTR exports from the endoplasmic reticulum (ER) to the Golgi apparatus and eventually reaches the plasma membrane, its molecular weight increases due to additional glycosylation (Figure S9).

**Figure 4.**
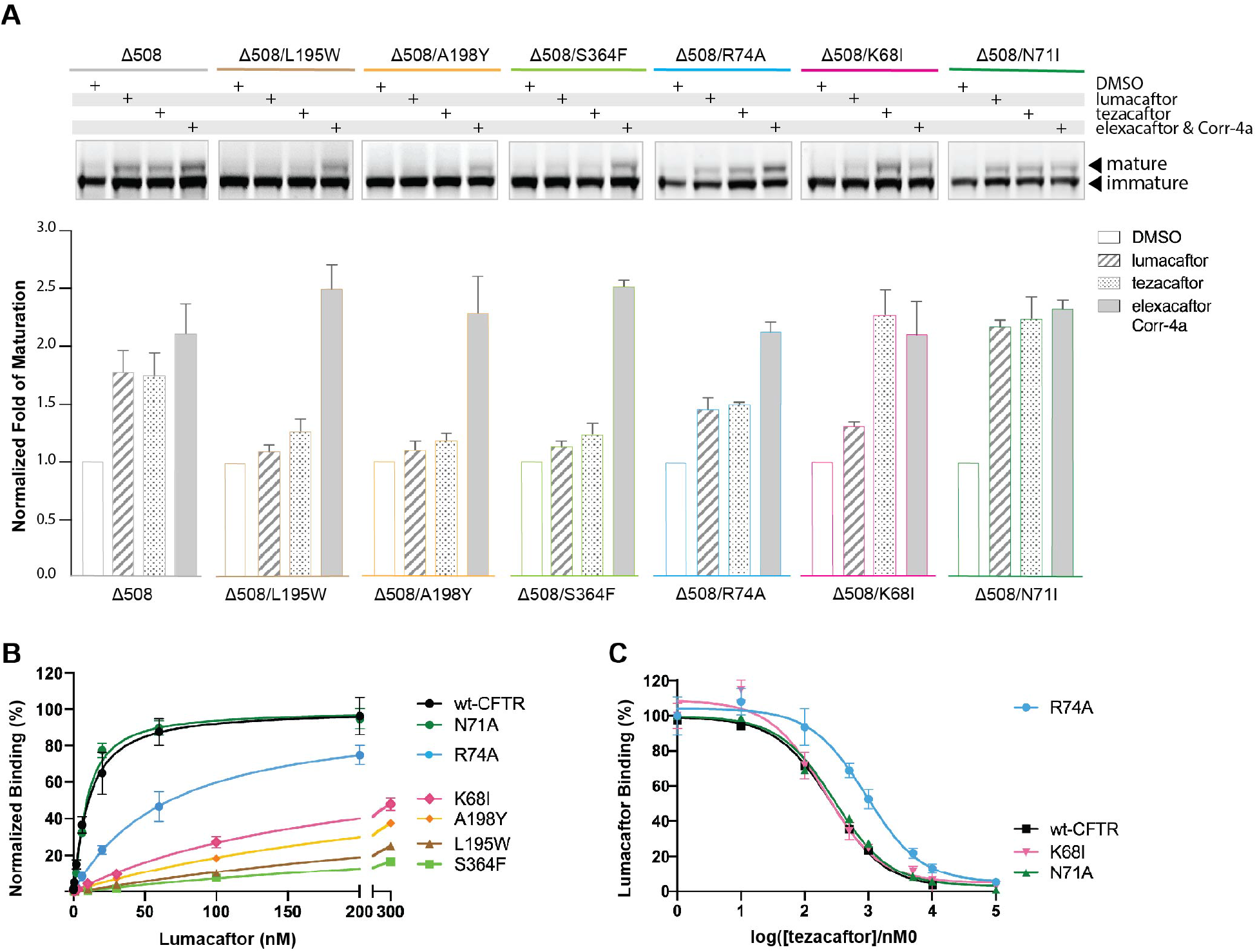
Mutations at the binding site diminished the efficacy of lumacaftor and tezacaftor. **(A)** Maturation assay of ΔF508-CFTR and binding-site mutations introduced to the ΔF508 background. Upper panel: SDS-PAGE of cell lysates from a single experiment, both mature and immature CFTR forms were visualized via the C-terminal GFP tag. Lower panel: Quantification of 3-6 repeats. The standard error of the mean is indicated as bars. Corrector concentrations: lumacaftor 1 μM, tezacaftor 10 μM, elexacaftor 0.2 μM, Corr-4a 10 μM in 0.1% DMSO. **(B)** SPA assay to measure the effects of mutations on lumacaftor binding. The K_d_ values of the polar residue substitutions K68I, R74A, and N71A were 0.19 ± 0.05 μM, 64 ± 13 nM, and 5.9 ± 2.9 nM, respectively. Those of the pocket-lining mutations A198Y, L195W, and S364F were 0.48 ± 0.11 μM, 0.86 ± 0.43 μM, and 1.43 ± 0.74 μM, respectively. Each data point represents the SEM of 6-9 of measurements. **(C)** Competition binding assay. The K_i_ values of K68I, N71A, and R74A CFTR were determined to be 0.12 ± 0.05 μM, 0.13 ± 0.03 μM and 0.41 ± 0.10 μM, respectively. For reference, the curves of the *wt*CFTR presented in Figure 1 were also shown (black line). Each data point represents the SEM of 6-9 of measurements. The concentration of [H^3^]lumacaftor was kept around the K_d_ value of the corresponding CFTR construct. See also Figures S8, S9.

Lumacaftor and tezacaftor increased the abundance of the mature, fully glycosylated form of the ΔF508 mutant (Figure 4A). This effect is severely diminished by the pocket-lining mutations L195W, A198Y, and S364F (Figure 4A). Furthermore, mutating the three polar residues R74, K68 and N71 generated different responses that are consistent with their structural roles in drug binding (Figure 4A). The R74A substitution lowers the efficacy of both lumacaftor and tezacaftor, consistent with the structures showing that it interacts with both drugs (Figure 2D, 3C-D and 3F-G). K68, on the other hand, forms a salt bridge with lumacaftor but makes no contact with tezacaftor (Figure 3D and 3F). Correspondingly, the K68I mutation diminished lumacaftor correction but did not affect tezacaftor. The side chain of N71 projects away from both drugs and its substitution did not affect either corrector (Figure 4A, 2D and 3C). None of the mutations had a significant effect on the efficacy of elexacaftor/Corr4a, drugs that belong to different classes of correctors (Figure 4A).

To further correlate the functional effects of the direct binding of lumacaftor, we mutated the same residues in the background of *wt*CFTR and measured their affinities for lumacaftor via SPA. The pocket-lining mutations severely reduced lumacaftor binding, likely due to steric-hindrance (Figure 4B). The K68I and R74A reduced the affinities by 23- and 8-fold, respectively (Figure 4B). In contrast, the N71A mutant exhibited an affinity similar to that of the *wt* protein (Figure 4B). The specificity of these perturbations is further demonstrated in the competition assay, which shows that the binding of tezacaftor is reduced by substitutions of R74, but not K68 nor N71 (Figure 4C). These effects on corrector affinities were not due to defects in folding, as these mutants behaved biochemically similarly to that of *wt* protein and showed strong binding to ivacaftor, a potentiator that interacts within the TMD2 of CFTR (Baroni et al., 2014; Liu et al., 2019) (Figure S10).

A recent study showed that lumacaftor does not rescue misfolded zebrafish CFTR (zCFTR) (Laselva et al., 2019) even though its overall structure is very similar to that of human CFTR (Liu et al., 2017; Zhang and Chen, 2016). Three amino acids distinguish the lumacaftor-binding site in these two CFTR orthologs. A pocket-lining methionine, corresponding to T360 in human CFTR, owing to its larger side chain, likely occludes lumacaftor binding in zCFTR. Furthermore, residues equivalent to K68 and R74 are serine and alanine in zCFTR, respectively. These differences are likely to diminish lumacaftor binding, resulting in its inability to rescue misfolded zCFTR.

## DISCUSSION

In this study, we present structures of CFTR in complex with type I correctors lumacaftor and tezacaftor. Consistent with the structure-activity relationship, these two analogous compounds bind to a common site in TMD1. The location of the binding site is entirely consistent with functional studies demonstrating that lumacaftor promotes folding of isolated TMD1 but does not affect other domains (Farinha et al., 2013; Kleizen et al., 2021; Laselva et al., 2018; Loo et al., 2013; Ren et al., 2013). Furthermore, binding of lumacaftor or tezacaftor did not alter the structure of CFTR, supporting the previous conclusion that lumacaftor stabilizes TMD1 in its native conformation (Loo et al., 2013). Binding-site mutations that lowered the affinities of lumacaftor and tezacaftor also diminished their abilities to rescue ΔF508-CFTR, indicating that the structurally identified type I corrector-binding site is the site of action of these drugs to restore CFTR folding.

Earlier work from Braakman and colleagues showed that individual domains of CFTR begin to adopt a tertiary structure as the nascent chain emerges from the ribosome (Kleizen et al., 2005). Folding is completed after the TMDs, NBDs, and R domain assemble into the final structure (Kleizen et al., 2005). The membrane-spanning region of CFTR contains a large number of polar residues, leading to inefficient and slow integration of TM helices in the membrane (Carlson et al., 2005; Hessa et al., 2005; Patrick et al., 2011). In addition, CFTR exhibits a domain-swapped configuration, such that TM helices 1, 2, 3 and 6 pack against TM 10-11 to form one bundle; and TM helices 4 and 5 interact with four helices in TMD2 to form another bundle. Such assembly cannot be established until the full-length protein is translated. It is estimated that CFTR synthesis takes about 10 minutes (Ward and Kopito, 1994) and the subsequent folding and assembly of TMDs and NBDs takes about 30-120 minutes (Amaral, 2004; Skach, 2006; Wang et al., 2006). During this process, partially folded intermediates linger in the ER, vulnerable to degradation. Consequently, only a small percentage of synthesized CFTR polypeptide reaches the cell surface even for the *wt* protein (Lukacs et al., 1994; Ward and Kopito, 1994). Folding mutations such as ΔF508 destabilize an individual domain and/or prevent effective interdomain assembly, leading to expansive pre-mature degradation (Cui et al., 2007; Davies et al., 2013; Du and Lukacs, 2009; Lukacs et al., 1994; Rosser et al., 2008; Serohijos et al., 2008; Younger et al., 2006).

The identification of the type I corrector binding site provides a structural basis to understand how these compounds promote CFTR folding. The N-terminal TMD1 is synthesized at an early stage of CFTR biogenesis and folds co-translationally (Kleizen et al., 2005). The 4 TM helices forming the corrector-binding site are predicted to be unstable. Using an established algorithm (Hessa et al., 2007) to calculate the free energy for membrane insertion (ΔG_insertion_), we find that TM helices 1, 2, 3, and 6 all have positive values, indicating that these helices individually are unstable in the membrane (Table S2). Instability was also confirmed experimentally for TM6 (Tector and Hartl, 1999). In addition, the tertiary structure formed by TM1, 2, 3 and 6 contains a hydrophobic pocket penetrating into the core of the protein (Figures 2B, 2F and 3A). Based on the classic work of Matthews, Bowie, and colleagues, the destabilizing energy caused by an internal cavity of this size (360 Å^3^) is substantial (Eriksson et al., 1992; Joh et al., 2009). Lumacaftor and tezacaftor, both largely hydrophobic and membrane-permeable, have negative ΔG values for membrane partitioning. Binding of these correctors would structurally link TM 1, 2, 3, and 6 together and contribute to a net negative ΔG for partitioning. In this manner, the type I correctors are able to stabilize the partially folded TMD1 while it awaits the completion of inter-domain assembly.

Consistent with this analysis, Clarke and colleagues showed that lumacaftor increased the lifetime of TMD1 by about 5-fold (Loo et al., 2013). And most recently, Braakman’s laboratory demonstrated that the type I correctors lumacaftor and C18 act at an early folding stage, supporting the hypothesis that rescuing ΔF508 by lumacaftor arises from the increased stability of TMD1 (Kleizen et al., 2021).

In summary, the aforementioned structural, theoretical, and experimental data collectively support the following mechanism of action for type I correctors (Figure 5). Once the N-terminal TMD1 is synthesized, it adopts a tertiary structure that is intrinsically unstable in the ER membrane. Binding of lumacaftor or tezacaftor stabilizes TMD1, making it less susceptible to targeted degradation by protein quality control machinery. As CFTR folding is a highly cooperative process, stabilizing TMD1 would ultimately increase the overall probability of forming a fully assembled structure and thereby allosterically rescue a large number of disease-causing mutants that reside in other parts of CFTR. This mechanism is also consistent with the synergy between lumacaftor and suppressor mutations (Farinha et al., 2013; Okiyoneda et al., 2013): lumacaftor extends the lifetime of TMD1 and the suppressor mutations stabilize different parts of CFTR or enhance inter-domain assembly, and thus together they achieve higher rescuing efficiency (Figure 5).

**Figure 5.**
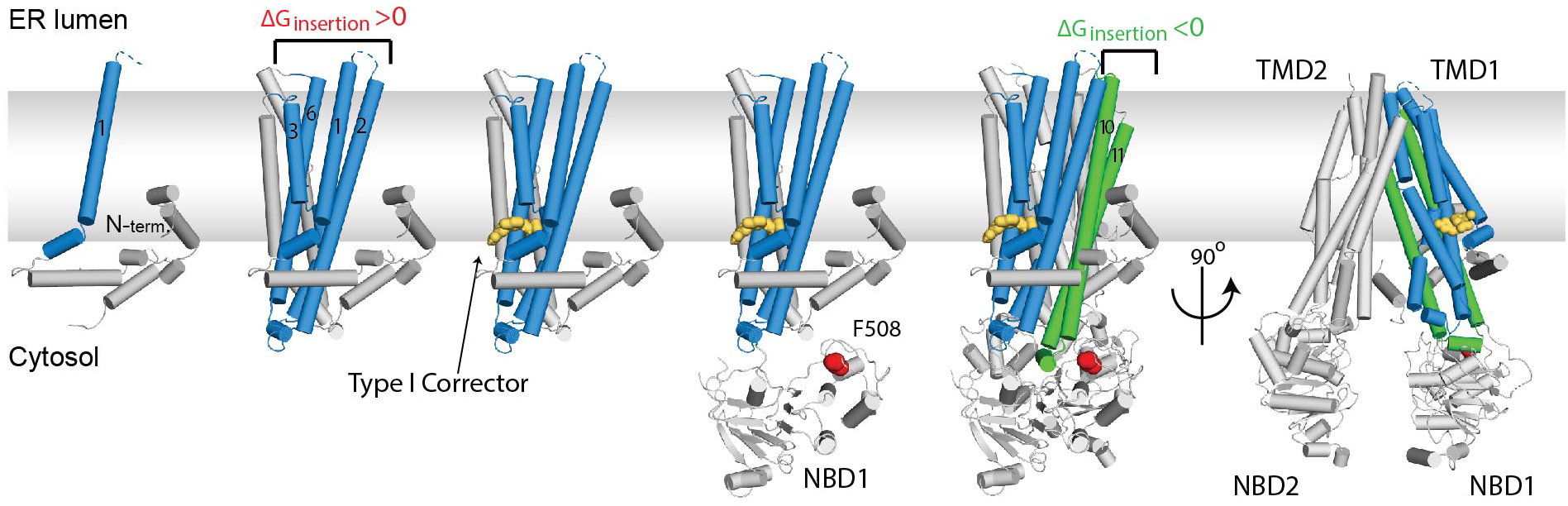
The proposed mechanism of type I correctors. CFTR folds co-translationally as individual domains are synthesized, followed by assembly of the mature tertiary structure. The N-terminal TMD1, synthesized in the early phase, is thermodynamically unstable. The binding of the corrector (yellow sticks) stabilizes TMD1 in the ER membrane, makes it less susceptible to degradation. Increasing the lifetime of TMD1 can partially rescue folding defects in other parts of CFTR, such as ΔF508 in NBD1 (indicated in red). For simplicity, the chaperones that assist CFTR folding are not shown. See also Table S2.

CFTR correctors, discovered empirically, are the most successful drugs to treat diseases caused by defects in protein folding. The proposed mechanism for CFTR correctors provides a conceptual framework to understand how a small molecule can influence protein folding. This concept, rooted in the energetics of protein folding, may also apply to other pharmacological chaperones targeting various misfolded proteins. As of today, most small molecule chaperones were developed as competitive inhibitors binding at the enzymatic active sites (Tran et al., 2020). The disadvantage of this approach is that the drug stabilizes folding of the disease-causing mutants but at the same time it diminishes enzymatic activity. An alternative approach would be to develop compounds that bind and increase the stability of an individual domain within the target protein. As most proteins fold co-translationally, such compounds can revert folding mutations through an allosteric effect as observed for the CFTR type I correctors.

## AUTHOR CONTRIBUTIONS

K.F. performed all the experiments. K.F. and J.C. conceptualized the study, analyzed the data, and wrote the manuscript. The authors declare no competing financial interests.

## ACKNOWLEDGMENTS

We thank D. Tallent for proofreading the manuscript, M. Ebrahim and J. Sotiris at Rockefeller’s Evelyn Gruss Lipper Cryo-Electron Microscopy Resource Center for assistance in data collection and F. Glickman of the Rockefeller High-throughput and Spectroscopy Resource Center for help with the SPA experiments. This work is supported by the Howard Hughes Medical Institute (to J.C) and the Cystic Fibrosis Foundation Therapeutics (to J.C and K.F).

---

**Figure S1.**
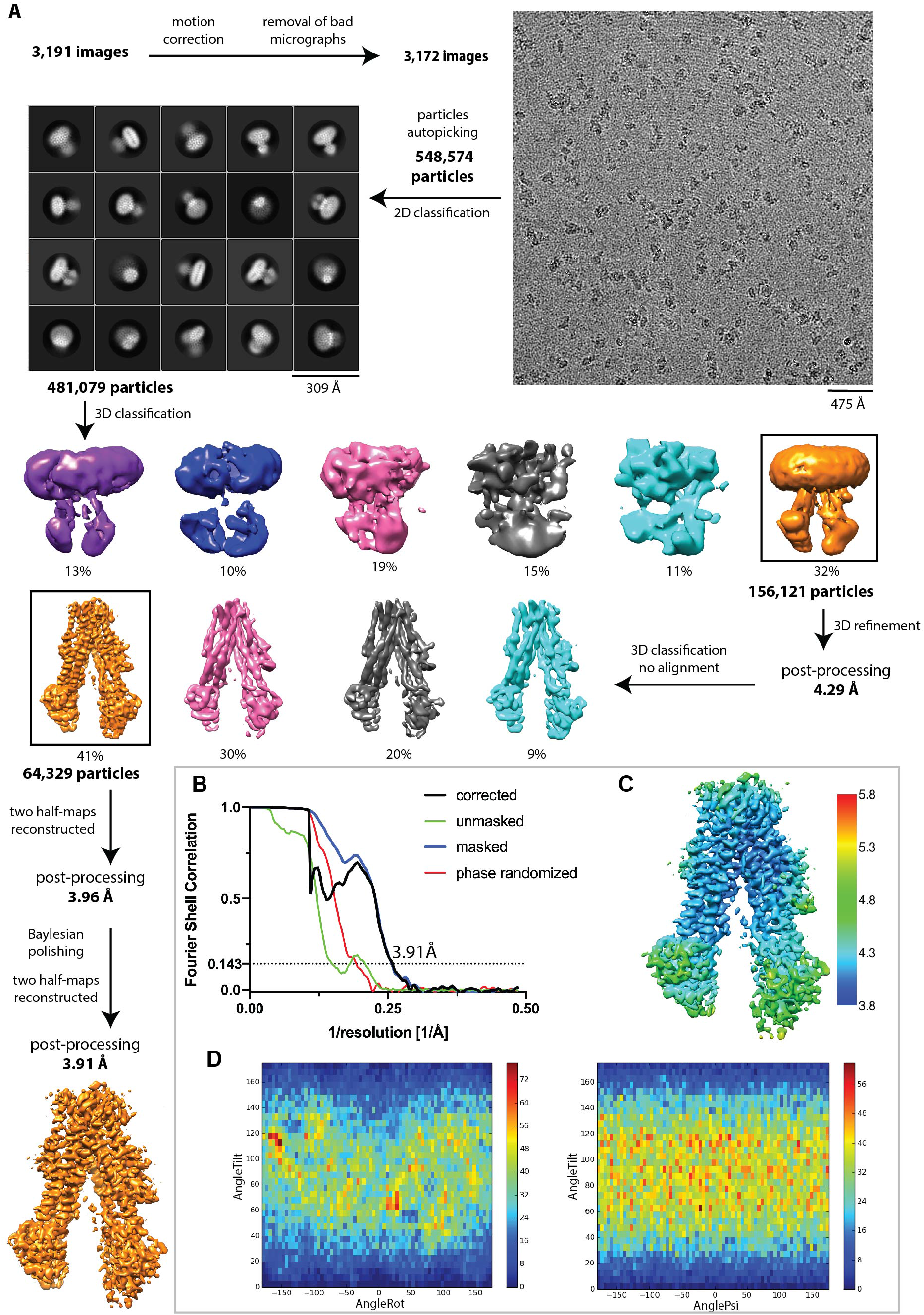
Cryo-EM analysis of the CFTR/lumacaftor complex. **(A)** Image processing procedure and representative example of a micrograph. **(B)** Fourier shell correlation curves of the final map. **(C)** Local resolution estimation of the final map. **(D)** Particles orientation distribution histograms. Related to Figure 2.

**Figure S2.**
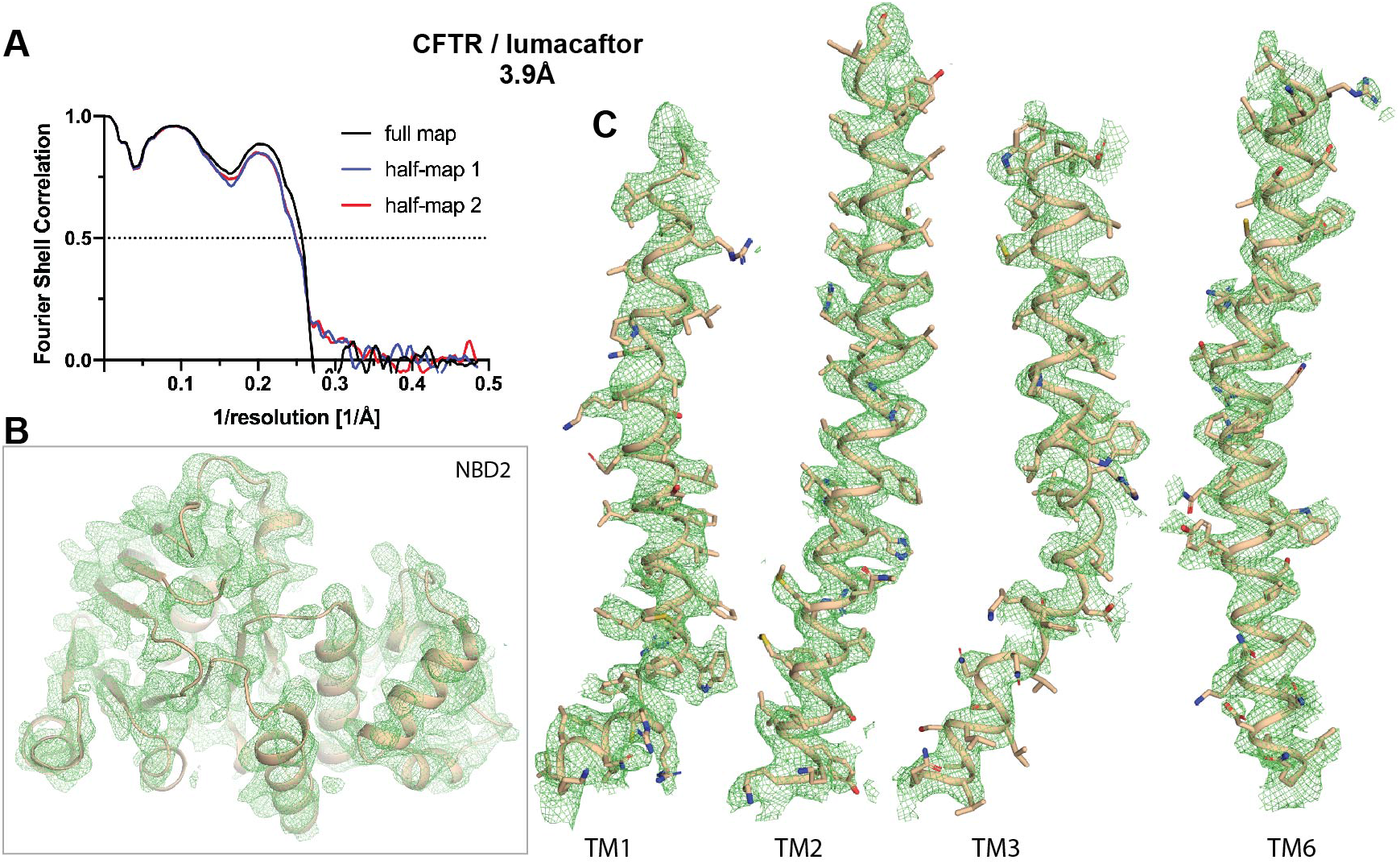
Quality of the CFTR/lumacaftor reconstruction. **(A)** Model-to-map fit for the full map (black), half-map 1 (blue), half-map 2 (red). **(B)** EM density of NBD2. **(C)** EM density of the four helices forming the lumacaftor-binding site. Related to Figure 2.

**Figure S3.**
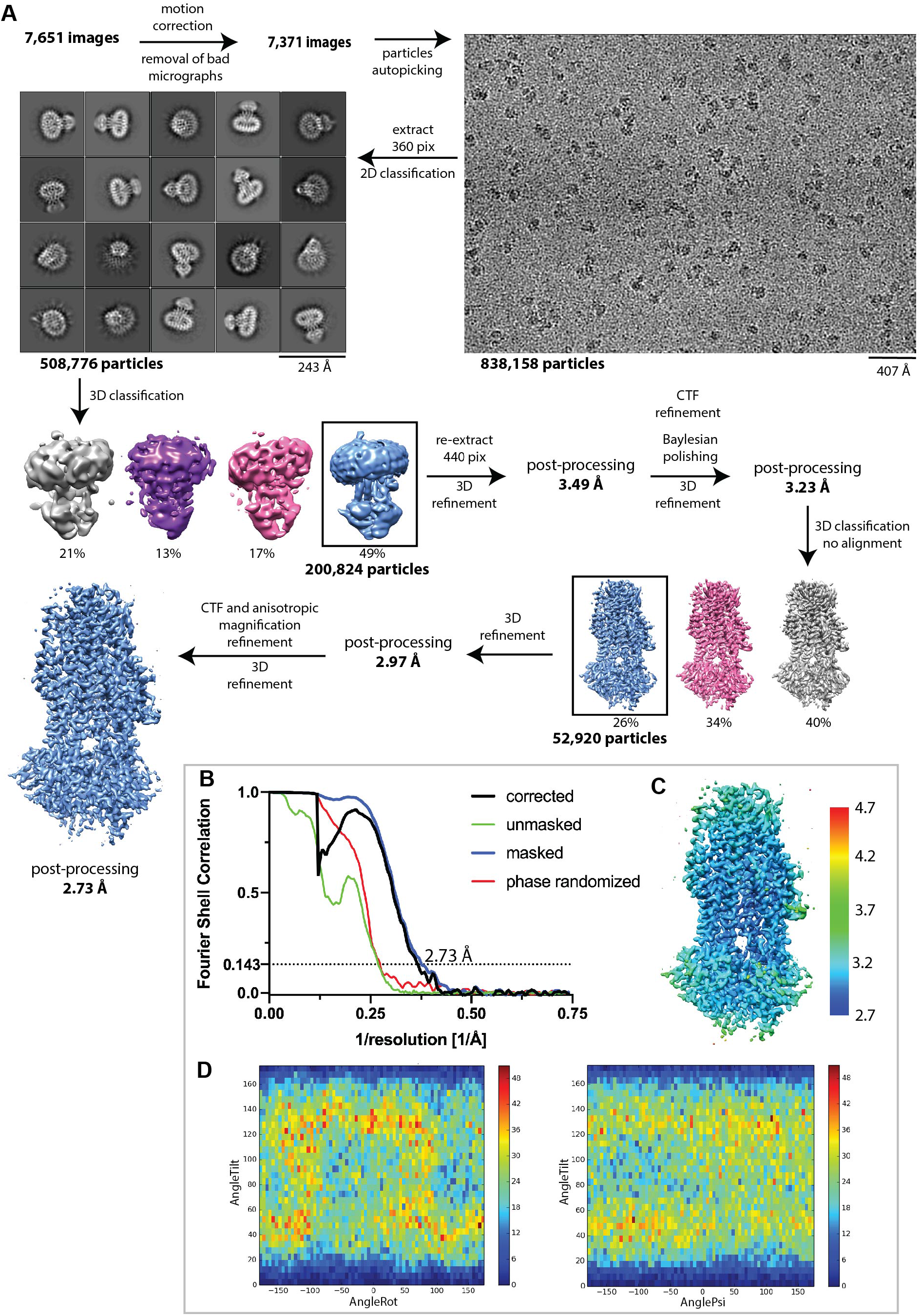
Cryo-EM analysis of the CFTR/lumacaftor+ATP complex. **(A)** Image processing procedure and representative example of a micrograph. **(B)** Fourier shell correlation curves of the final map. **(C)** Local resolution estimation of the final map. **(D)** Particles orientation distribution histograms. Related to Figure 2.

**Figure S4.**
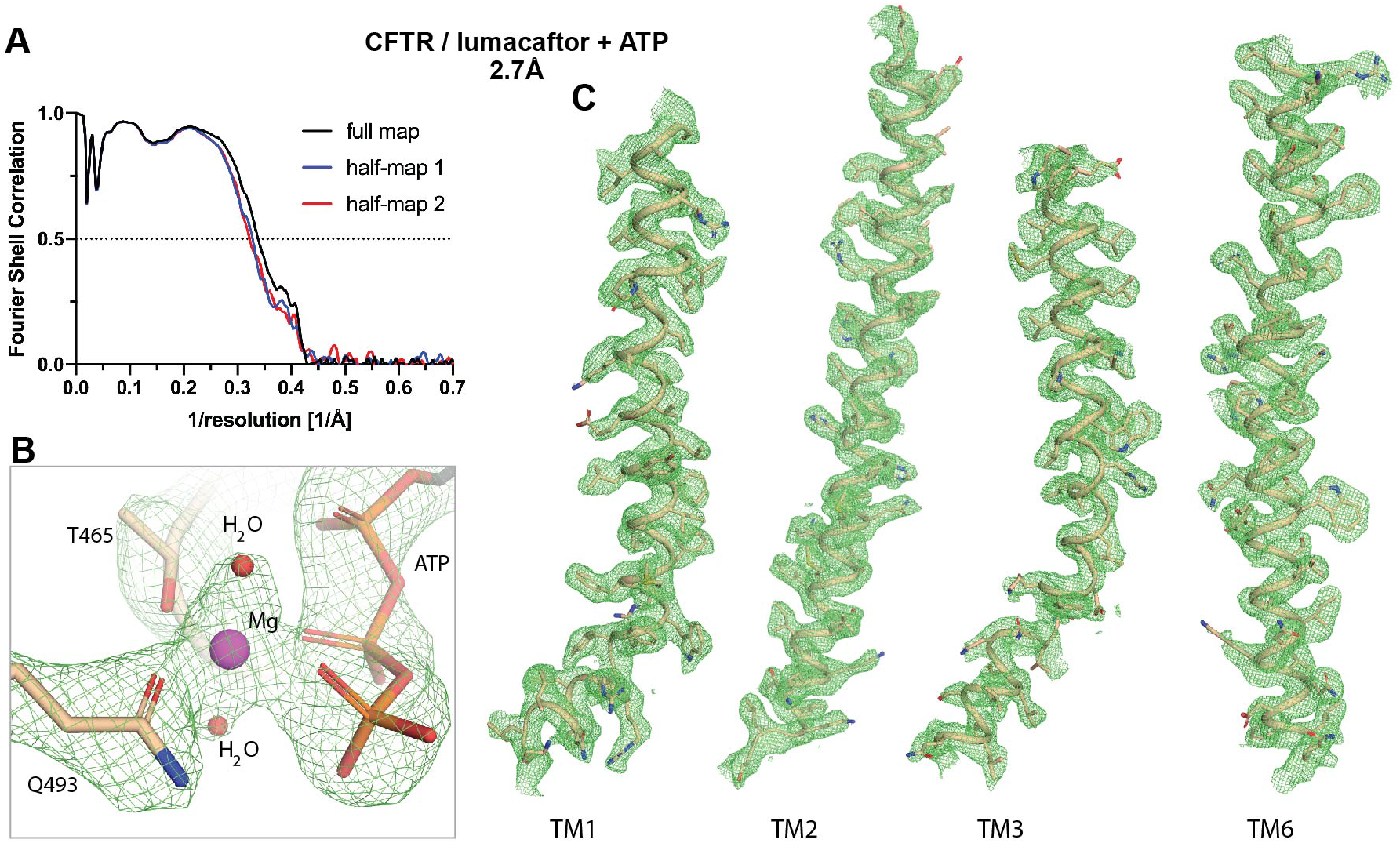
Quality of the CFTR/lumacaftor+ATP complex. **(A)** Model-to-map fit for the full map (black), halfmap 1 (blue), half-map 2 (red). **(B)** EM density at the degenerate ATP binding site. **(C)** EM density of the four helices forming the lumacaftor-binding site. Related to Figure 2.

**Figure S5.**
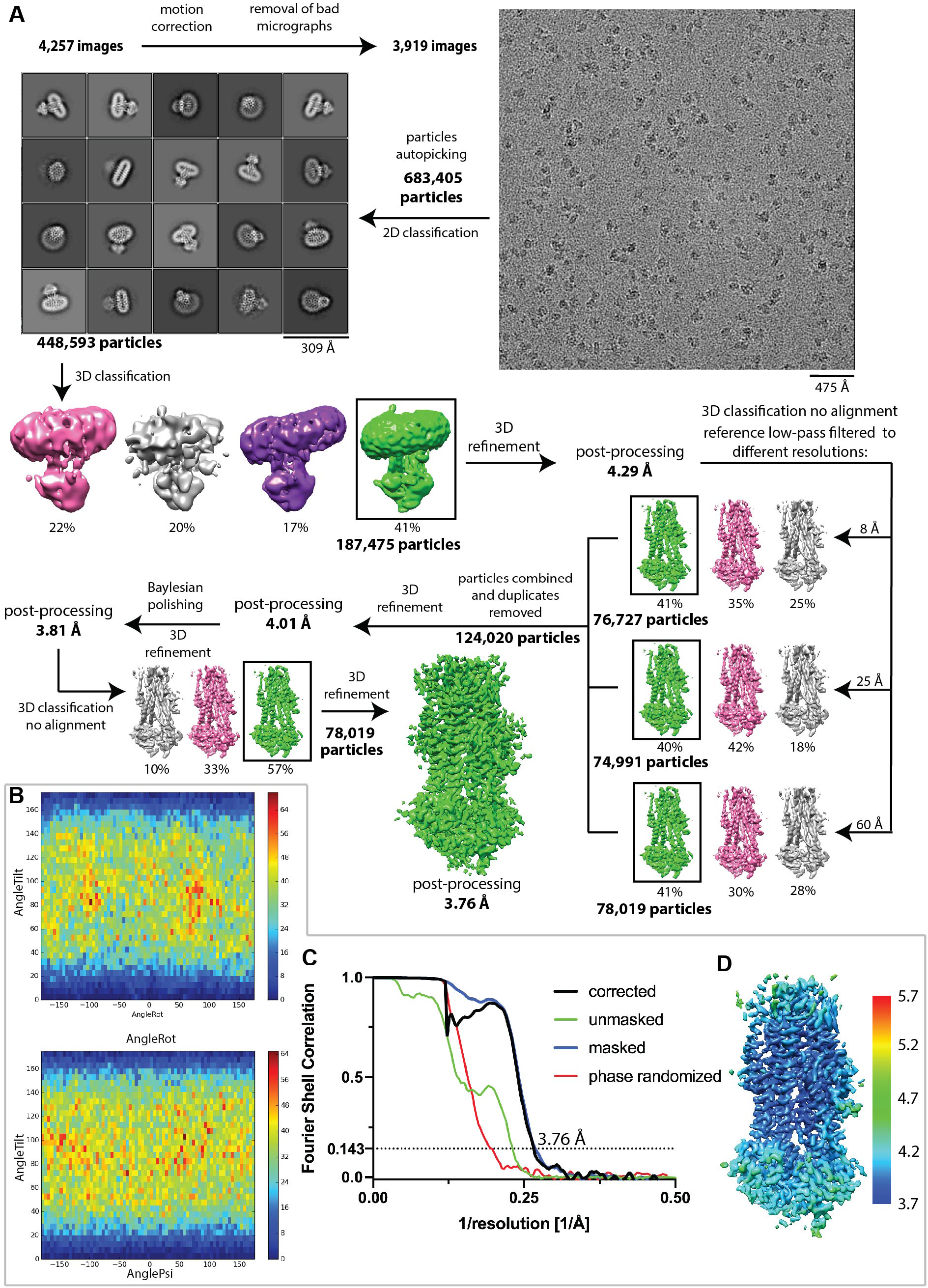
Cryo-EM analysis of the CFTR/tezacaftor+ATP complex. **(A)** Image processing procedure and representative example of a micrograph. **(B)** Particles orientation distribution histograms. **(C)** Fourier shell correlation curves of the final map. **(D)** Local resolution estimation of the final map. Related to Figure 3.

**Figure S6.**
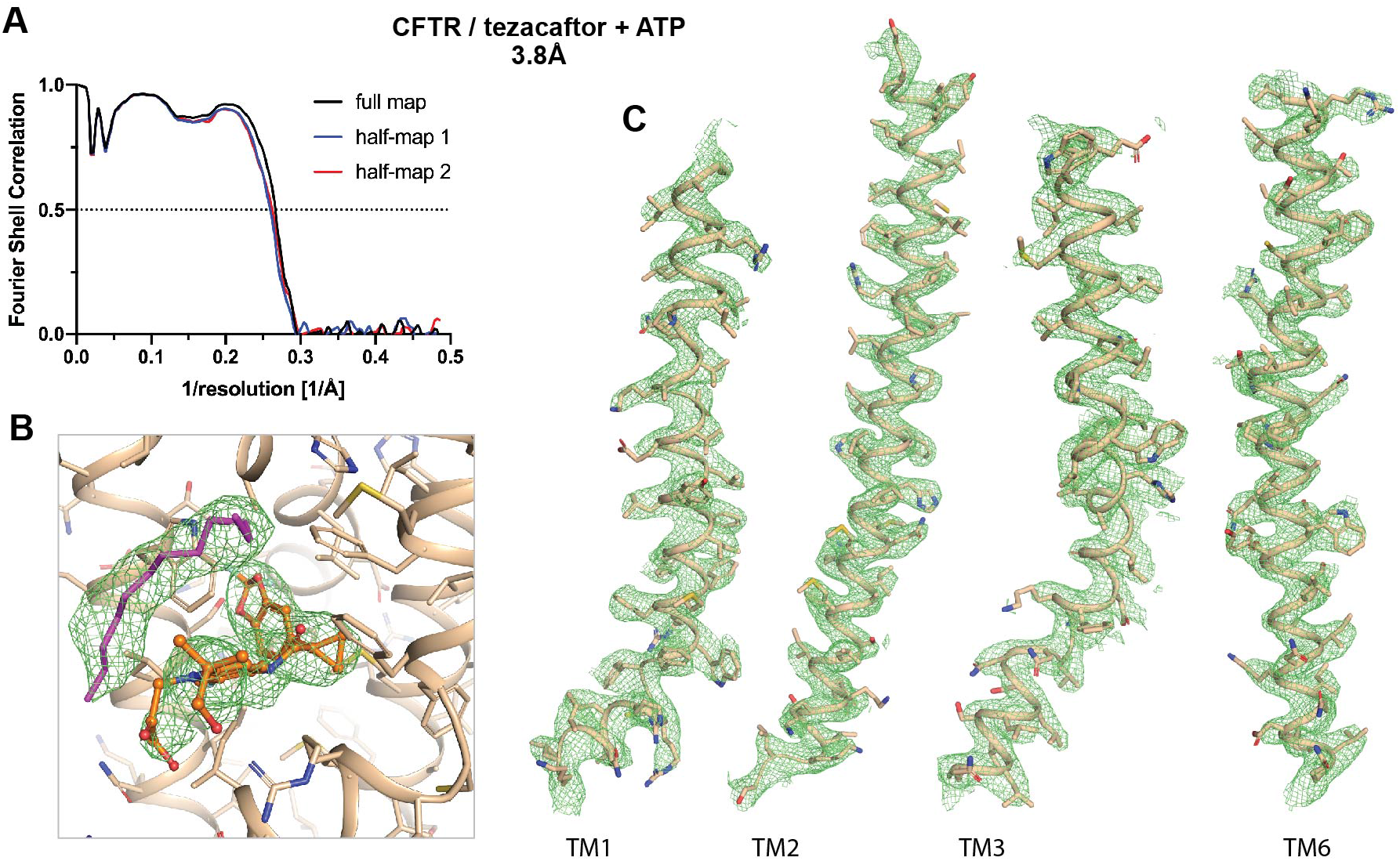
Quality of the CFTR/tezacaftor+ATP complex. **(A)** Model-to-map fit for the full map (black), halfmap 1 (blue), half-map 2 (red). **(B)** EM density at the tezacaftor-binding site. The lipid acyl chain is represented as magenta sticks. **(C)** EM density of the four helices forming the tezacaftor-binding site. Related to Figure 3.

**Figure S7.**
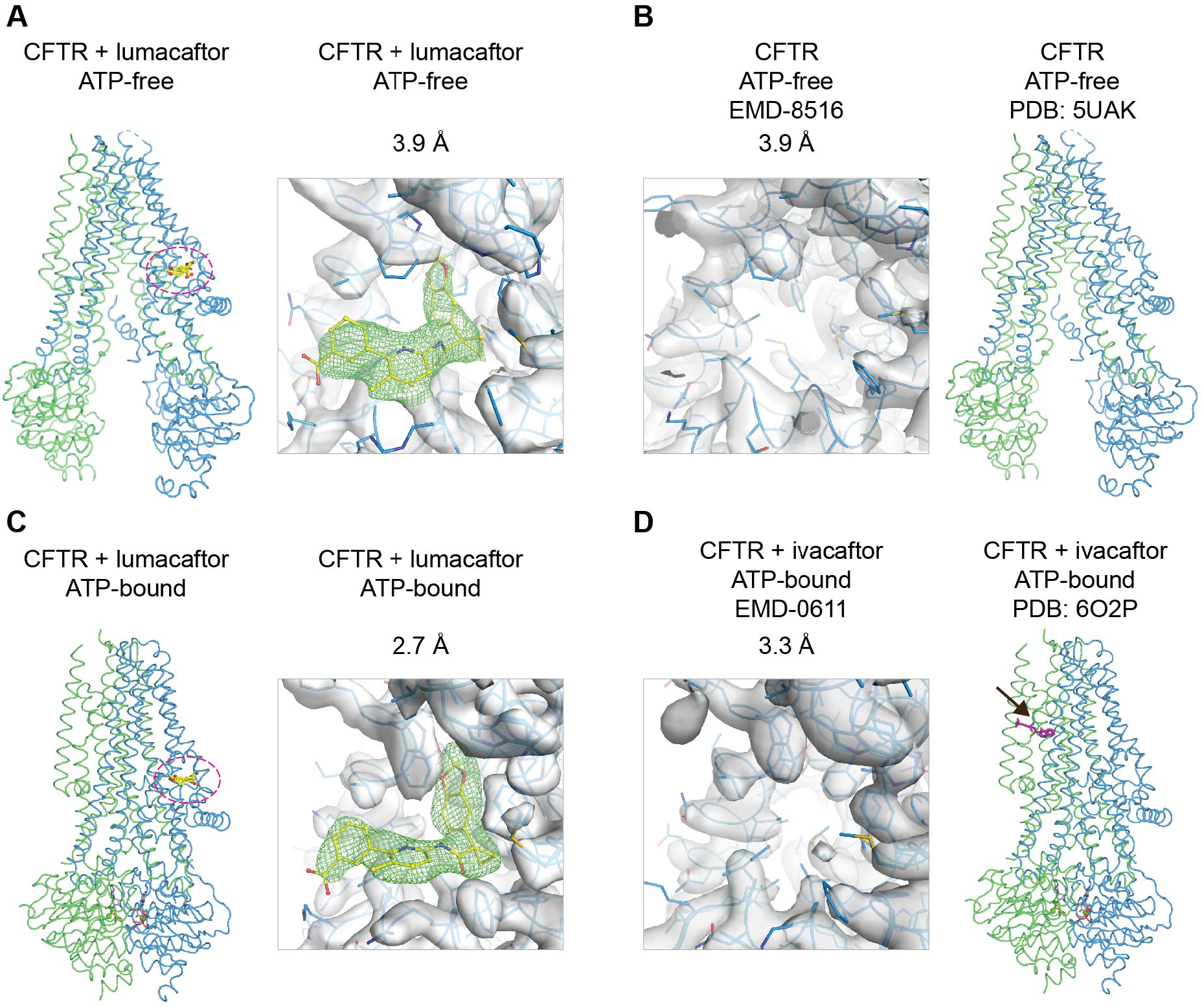
Comparison of the EM density at the corrector-binding site. **(A)** The lumacaftor-bound, NBD-separated structure (this study). **(B)** The apo structure (PDB:5UAK and EMD-8516). **(C)** The lumacaftor-bound, NBD-dimerized structure (this study). **(D)** The ivacaftor-bound structure (PDB:6O2P and EMD-0611). The black arrow indicates binding site of the potentiator ivacaftor (magenta sticks). All maps were contoured to show similar density for the CFTR main chain and side chains. Related to Figure 2.

**Figure S8.**
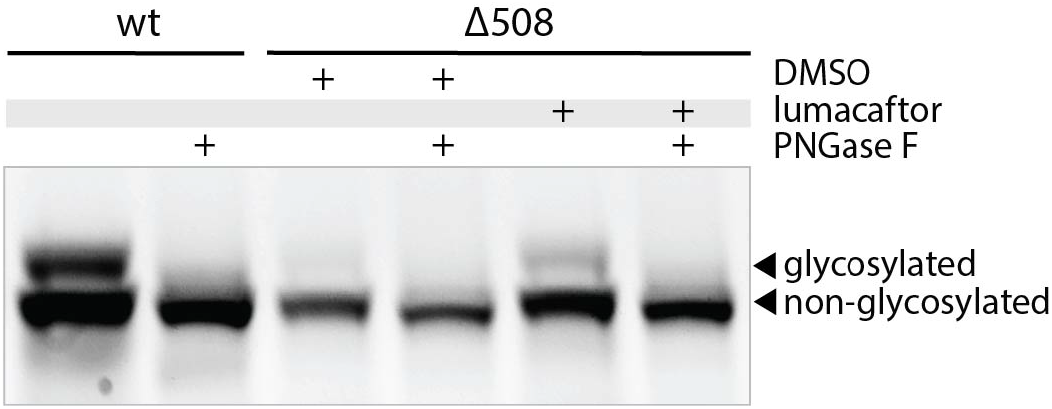
The mature, glycosylated form of CFTR is sensitive to PNGase F treatment. Related to Figure 4.

**Figure S9.**
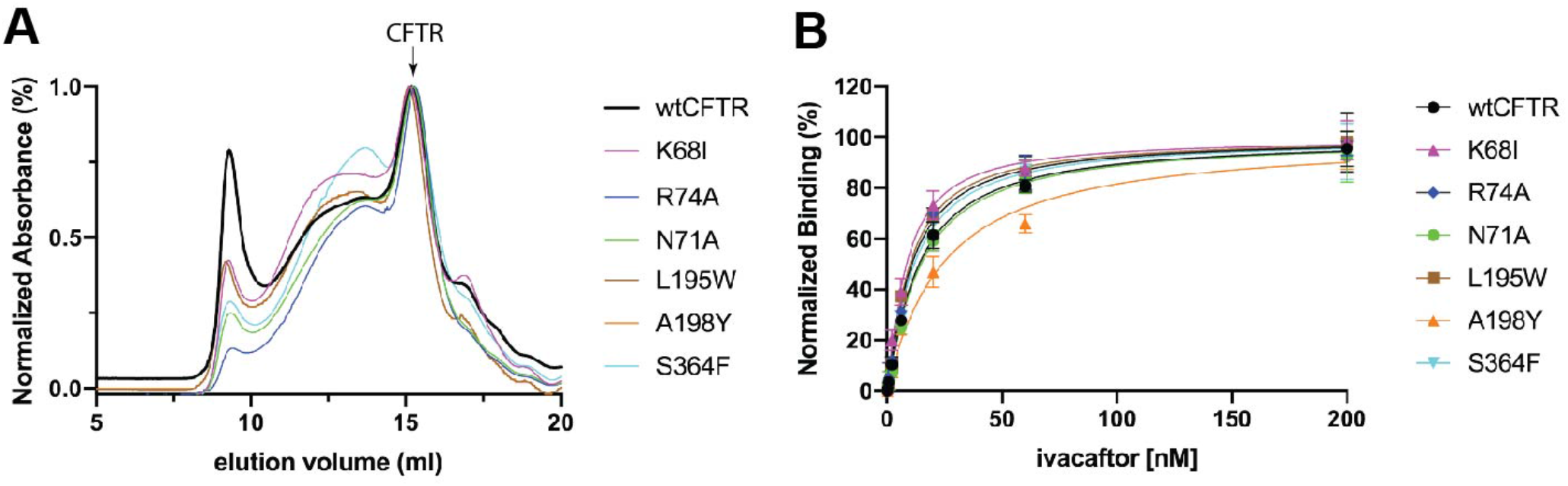
Folding and stability assessment of CFTR mutants used in this study. **(A)** Size exclusion chromatography profiles of the *wt* and mutant CFTR. The position of monomeric CFTR is indicated by an arrow. **(B)** Quantitative measurement of CFTR-ivacaftor interactions. The K_d_ values of the *wt*, K68I, R74A, N71A, L195W, A198Y and S364F CFTR were calculated to be 11.4 ± 2.5 nM, 6.1 ± 2.0 nM, 8.2 ± 2.0 nM, 12.1 ± 3.9 nM, 7.3 ± 2.6 nM, 21.9 ± 8.6 nM and 9.0 ± 2.6 nM respectively. Related to Figure 4.

**Figure S10.**
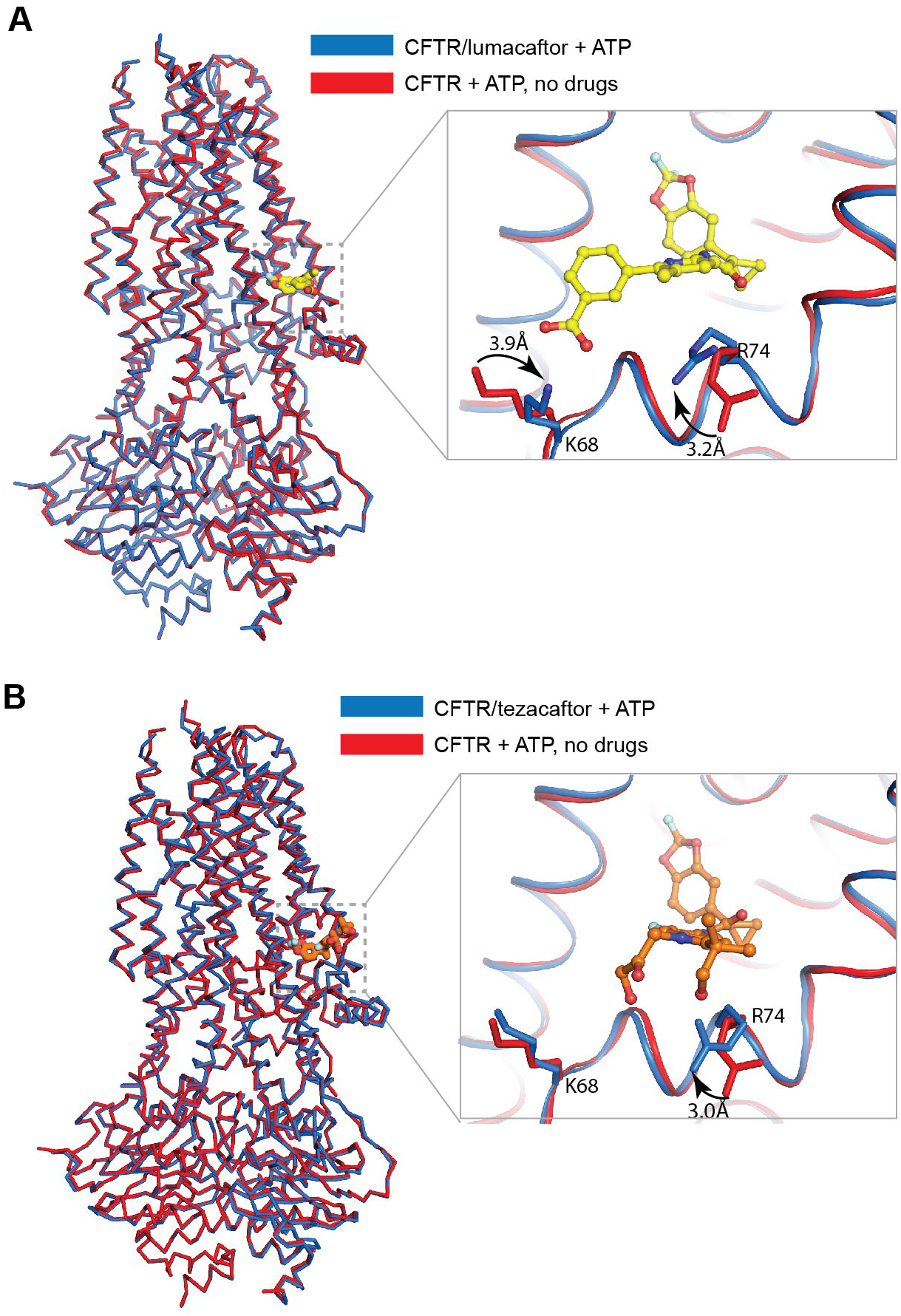
Comparison of the corrector-bound and drug-free CFTR structures. **(A)** Superposition of the CFTR/lumacaftor (blue) and drug-free (red) (PDB:6MSM) structures. Inset: local rearrangements of sidechains at the lumacaftor-binding site. Lumacaftor is represented in yellow sticks. **(B)** Superposition of the CFTR/tezacaftor (blue) and drug-free (red)(PDB:6MSM) structures. Inset: local rearrangements of sidechains the tezacaftor-binding site. Tezacaftor is represented in orange sticks. Related to Figure 2 and 3.

**Table S1.** Cryo-EM data collection, refinement and validation statistics.

|  | CFTR/lumacaftor -ATP<br>(EMDB-N/A)<br>(PDB N/A) | CFTR/lumacaftor +ATP<br>(EMDB- N/A)<br>(PDB N/A) | CFTR/tezacaftor +ATP<br>(EMDB- N/A)<br>(PDB N/A) |
| --- | --- | --- | --- |
| **Data collection and processing** |  |  |  |
| Magnification | 29,000 | 105,000 | 29,000 |
| Voltage (kV) | 300 | 300 | 300 |
| Electron exposure (e<sup>-</sup>/Å<sup>2</sup>) | 80 | 98 | 80 |
| Defocus range (μm) | 1.0 - 2.5 | 0.8 - 2.0 | 1.0 - 2.5 |
| Pixel size (Å) | 1.03 | 0.676 | 1.03 |
| Symmetry imposed | C1 | C1 | C1 |
| Initial particle images (no.) | 548,574 | 508,776 | 683,405 |
| Final particle images (no.) | 67,529 | 52,920 | 78,019 |
| Map resolution (Å) | 3.86 | 2.72 | 3.76 |
| FSC threshold | 0.143 | 0.143 | 0.143 |
| Map resolution range (Å) | 3.85 – 5.0 | 2.7 - 3.6 | 3.75 - 4.2 |
| **Refinement** |  |  |  |
| Initial model used (PDB code) | 5UAK | 6O1V | 6O1V |
| Model resolution (Å) | 3.9 | 2.7 | 3.8 |
| FSC threshold | 0.5 | 0.5 | 0.5 |
| Model resolution range (Å) | 3.85 – 5.0 | 2.7 - 3.6 | 3.75 - 4.2 |
| Map sharpening *B* factor (Å<sup>2</sup>) | -31.6 | -27.9 | -73.2 |
| Model composition |  |  |  |
| Non-hydrogen atoms | 7,841 | 9,635 | 9673 |
| Protein residues | 1,120 | 1,178 | 1,178 |
| Ligands | 1 | 12 | 14 |
| *B* factors (Å<sup>2</sup>) |  |  |  |
| Protein | 98 | 55 | 49 |
| Ligand (corrector) | 89 | 43 | 31 |
| R.m.s. deviations |  |  |  |
| Bond lengths (Å) | 0.003 | 0.004 | 0.005 |
| Bond angles (°) | 0.73 | 0.66 | 0.75 |
| Validation |  |  |  |
| MolProbity score | 1.67 | 1.36 | 1.41 |
| Clashscore | 6.79 | 4.01 | 5.70 |
| Poor rotamers (%) | 0 | 0 | 0 |
| Ramachandran plot |  |  |  |
| Favored (%) | 95.7 | 97.0 | 97.5 |
| Allowed (%) | 4.3 | 3.0 | 2.5 |
| Disallowed (%) | 0 | 0 | 0 |

**Table S2.**
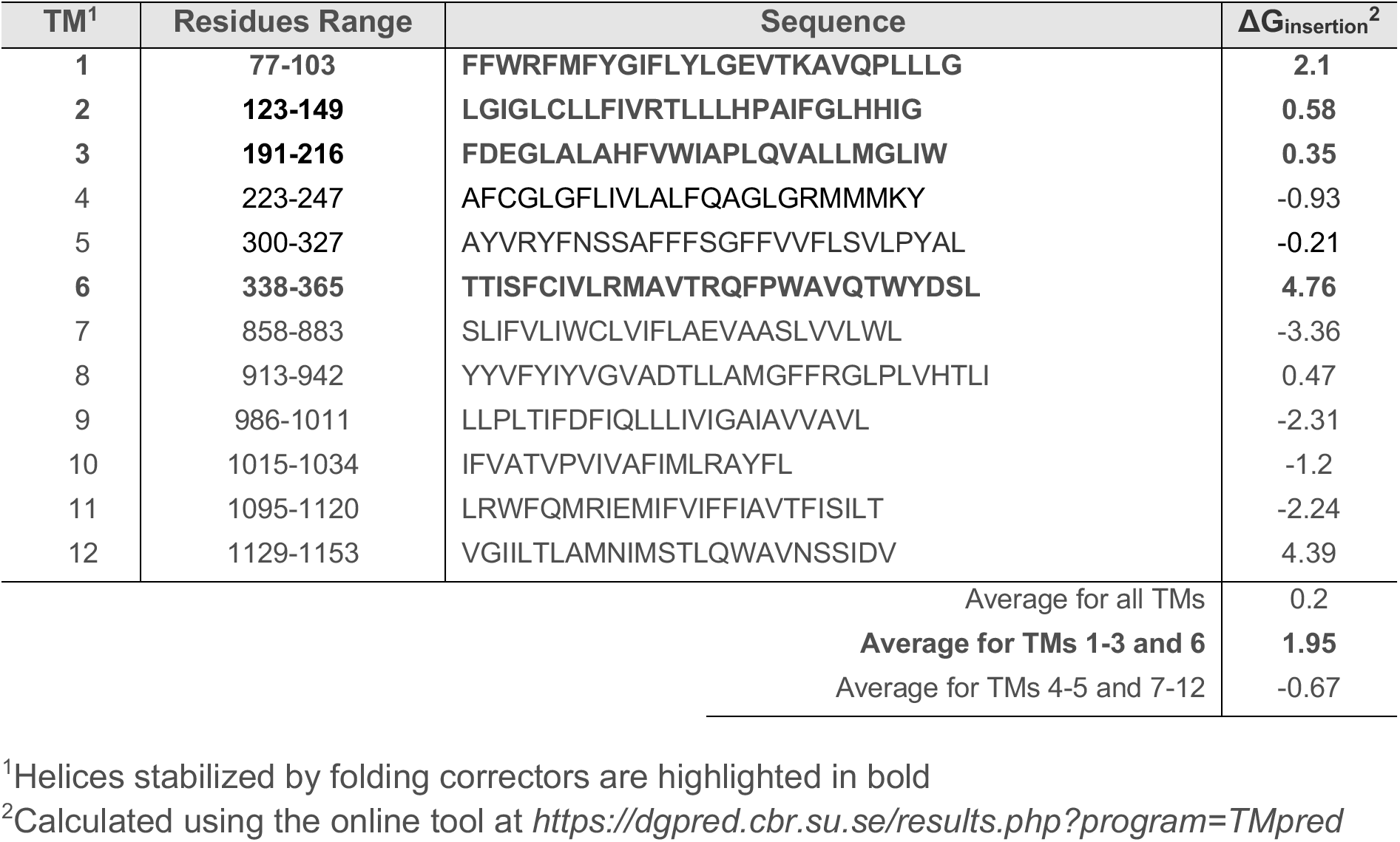
The predicted ΔG_insertion_ of the 12 TM helices.

## METHODS

### Cell culture

Sf9 cells were cultured in Sf-900 II SFM medium (GIBCO) supplemented with 5% FBS and 1% Antibiotic-Antimycotic. HEK293S GnTl^-^ cells were cultured in Freestyle 293 (GIBCO) supplemented with 2% FBS and 1% Antibiotic-Antimycotic. HEK293F cells were cultured in DMEM F-12 (ATCC) supplemented with 10% FBS and 1% Antibiotic-Antimycotic.

### Mutagenesis

All mutations were introduced using the SPRINP mutagenesis methodology (Edelheit et al., 2009).

### Protein expression and purification

All CFTR constructs were expressed and purified as described (Goehring et al., 2014; Zhang and Chen, 2016). Briefly, bacmids carrying CFTR constructs were generated in E. Coli DH10Bac cells (Invitrogen). Recombinant baculoviruses were produced and amplified in Sf9 cells. Proteins were expressed in HEK293S GnTl^-^ cells infected with 10% baculovirus at a density of 2.7×10^6^ cells/ml. Cells were induced with 10 mM sodium butyrate 12 hours after infection and cultured at 30° C for another 48 hours before harvesting. For protein purification, cells were solubilized in buffer containing 1.2% 2,2-didecylpropane-1,3-bis-β-D-maltopyranoside (LMNG) and 0.24% Cholesteryl hemisuccinate (CHS). Protein was purified via its C-terminal green fluorescence protein (GFP) tag using GFP nanobody coupled Sepharose Beads (GE Healthcare) and eluted by removing the GFP tag with the PreScission Protease. The E1371Q samples were phosphorylated using protein kinase A (NEB). The wild-type sample was de-phosphorylated using λ-phosphatase. The CFTR (E1371Q) sample was phosphorylated with protein kinase A. At the final step, protein samples were purified on size exclusion chromatography in 0.06% (wtCFTR/lumacaftor and CFTR (E1371Q)/tezacaftor samples) or 0.03% (CFTR (E1371Q)/lumacaftor) digitonin.

### EM data acquisition and processing

Immediately after size exclusion chromatography, the CFTR (E1371Q) sample (at 5 mg/mL) was incubated with 10 mM ATP, 8 mM MgCl_2_ and 200 *μ*M lumacaftor or tezacaftor on ice for 30 min. The *wt*CFTR sample (5mg/ml; 32 *μ*M) was incubated with 200 *μ*M lumacaftor. About 3 mM fluorinated Fos-choline-8 was added to the samples right before freezing on to Quantifoil R1.2/1.3 400 mesh Au grids using Vitrobot Mark IV (FEI).

Cryo-EM images were collected on a 300 kV Titian Krios (FEI) with a K2 or K3 Summit detector (Gatan) using SerialEM (Table S1). The images were corrected for gain reference and binned by 2. Drift correction was performed using MotionCorr (Zheng et al., 2017). Contrast transfer function (CTF) estimation was performed using CTFFIND4 (Rohou and Grigorieff, 2015) and GCTF (Zhang, 2016). Based on CTFFIND4 results, all the images at resolution lower than 5Å were removed. For further processing steps, GCTF generated values were used. Particles were automatically picked by Gautomatch (http://www.mrc-lmb.cam.ac.uk/kzhang/) for the *wt*CFTR/lumacaftor and CFTR(E1371Q)/tezacaftor datasets. For the CFTR (E1371Q)/lumacaftor data, picking was performed using RELION implemented Laplacian-of-Gaussian blob detection. All the subsequent steps of maps reconstruction and resolution estimations were performed using RELION 3.1(Scheres, 2012) (Figure S1, S3, S5).

### Model building and refinement

The initial protein models were built by fitting the published CFTR structures (PDB:5UAK and 6O1V) into the cryo-EM maps using UCSF Chimera (Pettersen et al., 2004). In the wtCFTR/lumacaftor structure, the sidechains of the NBDs were trimmed due to the limited resolution of ~4.5 Å (Figure S2). Models were then adjusted based on the cryo-EM densities using Coot (Emsley and Cowtan, 2004). Lumacaftor and tezacaftor were built into the drug density and refined in PHENIX (Adams et al., 2010) using restrains generated by eLBOW (Moriarty et al., 2009). MolProbity (Chen et al., 2010) was used for geometries validation.

Model overfitting was assessed as described (Johnson and Chen, 2017). Each model refined against half-map 1, converted to an electron density map using UCSF Chimera, and SPIDER (Shaikh et al., 2008) was used to calculate FSC plots between the converted map and the full map, the half-map 1, and the half-map 2. The cryo-EM maps were masked with a generous mask about 3.5 times larger than the volume of the model density. The FSC plots were then corrected for the volume by which the mask exceeds the volume of the model density using equation 1:

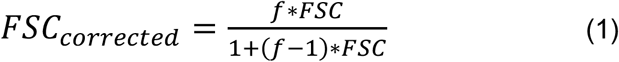

Where *f* is equal to the factor by which the mask exceeds the volume of the model density (Figure S2, S4, S6).

### Maturation assay

HEK293F cells grown in a 6-well plate were transiently transfected with CFTR constructs labelled with C-terminal eGFP tag using Lipofectamine 3000 (Thermo Fisher) in Opti-MEM (GIBCO) medium. Cells were incubated with DNA/transfection mixture for 12 hours at 37° C, then in DMEM F-12 supplemented with 10 mM sodium butyrate and the corrector of choice at 30° C for another 24 hours. Cells were harvested by re-suspending in 1 mL ice-cold PBS and spun down in 1.5 mL tubes for 5 min. at 5,000 rpm,4° C.

Cell pellets were re-suspended in buffer containing 1.2% 2,2-didecylpropane-1,3-bis-β-D-maltopyranoside (LMNG) and 0.24% Cholesteryl hemisuccinate (CHS) and rotated for 60 min. at 4° C. Cell lysates were spun down for 60 min. at 45,000 rpm, supernatants were analyzed on a 4-20% gradient tris-glycine SDS-PAGE gel (Thermo Fisher). Gels were imaged to visualize the GFP signal, which was quantified using Fiji (Schindelin et al., 2012). The background signal was subtracted from the CFTR bands. The proportion of mature CFTR to total CFTR (k_m/t_) was calculated using equation 2 and then normalized to that of the DMSO treated sample.

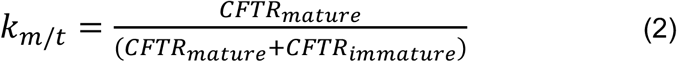

### Scintillation proximity assay

The binding and competition assays were performed as described (Liu et al., 2019). CFTR constructs used in this assay contain a C-terminal Strep-tag, followed by a PreScission Protease cleavage site, and a GFP tag. The GFP tag was removed during purification whereas the Strep-tag was retained to attach CFTR to the SPA beads. To measure lumacaftor binding, 5 nM CFTR was incubated with 0.5 mg/ml YSi streptavidin SPA beads (PerkinElmer) in the presence of varying concentrations of lumacaftor (at 1:1 molar ratio of cold and [^3^H] lumacaftor (6.4 Ci/mmol, synthesized by Moravek) in buffer containing 20 mM Tris-HCl pH 7.5, 200 mM NaCl, 0.06% digitonin, 2 mM DTT and 0.1% Tween 20 at 4° C for 1 hr. The reactions were carried out in 96-well non-binding surface microplates (Corning) and data were recorded using a Microbeta Trilux plate reader (PerkinElmer). Specific binding was obtained by subtracting background radioligand binding in the absence of protein. The K_d_ values were calculated by fitting the data with a single-site saturation binding model accounting for ligand depletion using GraphPad Prism 8 (GraphPad Software, San Diego, California, USA, www.graphpad.com). The K_i_ values of tezacaftor were calculated by fitting the data with a single-site competitive binding model in GraphPad Prism 8. The readings were normalized by dividing the specific binding with the total binding (Bmax) and represented in percentages.

### Data Presentation

Structural figures were generated using UCSF Chimera, PyMOL and Fiji. Plots were generated using GraphPad Prism8. Particles Euler angle histograms were generated using *plot_indiveuler_histogram_fromstarfile.py* script (https://github.com/leschzinerlab/Relion). All the figures were assembled using Adobe Illustrator.

